# δ-protocadherins control neural progenitor cell proliferation by antagonizing Ryk and Wnt/β-catenin signaling

**DOI:** 10.1101/2020.09.14.297283

**Authors:** Sayantanee Biswas, Michelle R. Emond, Kurtis Chenoweth, James D. Jontes

## Abstract

The proliferation of neural progenitor cells provides the cellular substrate from which the nervous system is sculpted during development. The δ-protocadherin family of homophilic cell adhesion molecules is essential for the normal development of the nervous system and has been linked to an array of neurodevelopmental disorders. However, the biological functions of δ-protocadherins are not well-defined. Here, we show that the δ-protocadherins regulate proliferation in neural progenitor cells, as lesions in each of six, individual δ-protocadherin genes increase cell division in the developing hindbrain. Moreover, Wnt/β-catenin signaling is upregulated in δ-protocadherin mutants and inhibition of the canonical Wnt pathway occludes the observed proliferation increases. We show that the δ-protocadherins physically associate with the Wnt receptor Ryk, and that Ryk is required for the increased proliferation in protocadherin mutants. Thus, the δ-protocadherins act as novel regulators of Wnt/β-catenin signaling during neural development and could provide lineage-restricted local regulation of canonical Wnt signaling and cell proliferation.

## Introduction

The regulated production of new cells through proliferative cell divisions is one of the primary driving forces in the development of multicellular organisms. The correct number of new cells and cell types must be temporally and spatially regulated during development. The canonical Wnt/β-catenin pathway plays prominent roles in regulating cell proliferation during development, and mutations in components of this pathway contribute to cancers and neurodevelopmental disorders (Cadigan and Liu, 2006; Clevers, 2006; De Ferrari and Moon, 2006; Kwan et al., 2016; MacDonald et al., 2009; Nusse and Clevers, 2017). However, the mechanisms providing spatiotemporal context to the regulation of proliferation and Wnt/β-catenin signaling are less well understood.

The δ-protocadherins (δ-pcdhs) are homophilic cell adhesion molecules within the cadherin superfamily (Bisogni et al., 2018; Cooper et al., 2016; Harrison et al., 2020), that are subdivided into δ1 and δ2 subfamilies on the basis of the number of extracellular cadherin repeats (7 for δ1-pcdhs and 6 for δ2-pcdhs) and conserved sequence motifs in their intracellular domains (Hulpiau and van Roy, 2011; Vanhalst et al., 2005). In the zebrafish, there are five δ1-pcdhs (*pcdh1a, pcdh1b, pcdh7a, pcdh7b* and *pcdh9*) and six δ2-pcdhs (*pcdh10a, pcdh10b, pcdh17, pcdh18a, pcdh18b* and *pcdh19*), as well as the δ2-like paraxial protocadherin. The δ-pcdhs are essential for neural development, as they have been implicated in a wide range of neurodevelopmental disorders (Hirano and Takeichi, 2012; Redies et al., 2012), including autism spectrum disorders (Bruining et al., 2015; Morrow et al., 2008; Piton et al., 2011) and epilepsy (Depienne et al., 2009; Dibbens et al., 2008; Lal et al., 2015; Perez-Palma et al., 2017). Mutations in the δ2-pcdh, *PCDH19*, result in a female-limited form of infant-onset epilepsy (Depienne et al., 2009; Dibbens et al., 2008), making *PCDH19* the second most commonly affected gene in epilepsies (Depienne and LeGuern, 2012). As well as their importance for brain development, the δ-pcdhs are intimately associated with cancer (Berx and van Roy, 2009; van Roy, 2014). Expression of δ-pcdhs is downregulated in multiple cancer cell lines with various examples of silencing by promoter methylation (Tang et al., 2012; Zhang et al., 2017). In addition, prognosis correlates with levels of δ-pcdh expression (Bing et al., 2018), while invasion and metastasis correlate with δ-pcdh loss (Lin et al., 2018). Additionally, there is evidence that expression of δ-pcdhs can inhibit tumor growth through regulating Wnt/β-catenin signaling (Xu et al., 2015; Yin et al., 2016; Zong et al., 2017). It was also reported that the clustered protocadherin, γ-C3 isoform, can interact with Axin to inhibit canonical Wnt signaling (Mah et al., 2016).

Members of the δ-pcdh family fulfill a variety of roles during development, including the regulation of morphogenetic cell movements (Aamar and Dawid, 2008; Biswas et al., 2010; Chen and Gumbiner, 2006; Emond et al., 2009; Kim et al., 1998; Kraft et al., 2012; Williams et al., 2018), axon outgrowth and guidance (Biswas et al., 2014a; Hayashi et al., 2014; Leung et al., 2013; Piper et al., 2008) and dendrite morphogenesis (Wu et al., 2015). Here, we show that in zebrafish, the δ-pcdh family regulates cell proliferation in the developing neuroepithelium, suggesting that regulation of cell division is a core function for this family of proteins. In addition, we provide evidence that δ-pcdhs influence cell proliferation by regulating Wnt/β-catenin signaling, as the canonical Wnt pathway is elevated in each of the δ-pcdh mutants, and inhibiting this pathway blocks the elevated cell proliferation observed in these mutants. We further show that the Wnt receptor Ryk physically interacts with the δ-pcdhs and is required for the increased proliferation. These results suggest that the δ-pcdhs are novel upstream regulators of canonical Wnt/β-catenin signaling, and provides a link between their biological role and their involvement in neurodevelopmental disorders and cancers.

## Results and Discussion

We previously showed that zebrafish *pcdh19* is expressed in neural progenitor cells in the zebrafish optic tectum and that Pcdh19 regulates proliferation. Mutant embryos lacking *pcdh19* exhibited an increased proliferation, as assessed by EdU labeling and phospho-Histone H3 (pH3) immunostaining (Cooper et al., 2015). The increased proliferation was accompanied by an increased number of *pcdh19*+ neurons. Similar results have been obtained for mouse *pcdh11x*, which negatively regulates proliferation and neurogenesis in mouse cortex (Zhang et al., 2014). Pcdh19 and Pcdh11x are members of the δ-pcdh family, δ2- and δ1-pcdh subfamilies, respectively (**Figure 1A**). To determine if other members of the δ-pcdh family play a similar role, we used CRISPR/Cas9 to make targeted lesions in the extracellular domains of *pcdh1a, pcdh7a, pcdh9* and *pcdh17* (**Supplemental Figure 1**). We used TALENs to generate mutations in *pcdh18b* and previously reported TALEN-mediated lesions in *pcdh19* (Cooper et al., 2015). Thus, we generated mutant lines for three of five δ1-pcdhs and three of six δ2-pcdhs. Like *pcdh19*, each of the other δ-pcdhs is expressed in the neuroepithelium at this stage of development and at later stages (**Supplemental Figure 2**). To assess the roles of these δ-pcdhs in regulating proliferation, we performed pH3 immunostaining, and counted the total number of pH3+ cells in the hindbrains of 18 hours post-fertilization (hpf) embryos (**Figure 1B-D**). As shown previously in the optic tectum, loss of *pcdh19* resulted in an increase in pH3+ cells in the hindbrain, which was rescued by forced expression of Pcdh19-GFP (**Figure 1E**). For each δ-pcdh that we tested, we also observed an elevation in cell proliferation (**Figure 1F**). These results suggest that a central function of the δ-pcdh family is to regulate proliferation in neural progenitor cells.

**Figure 1.**
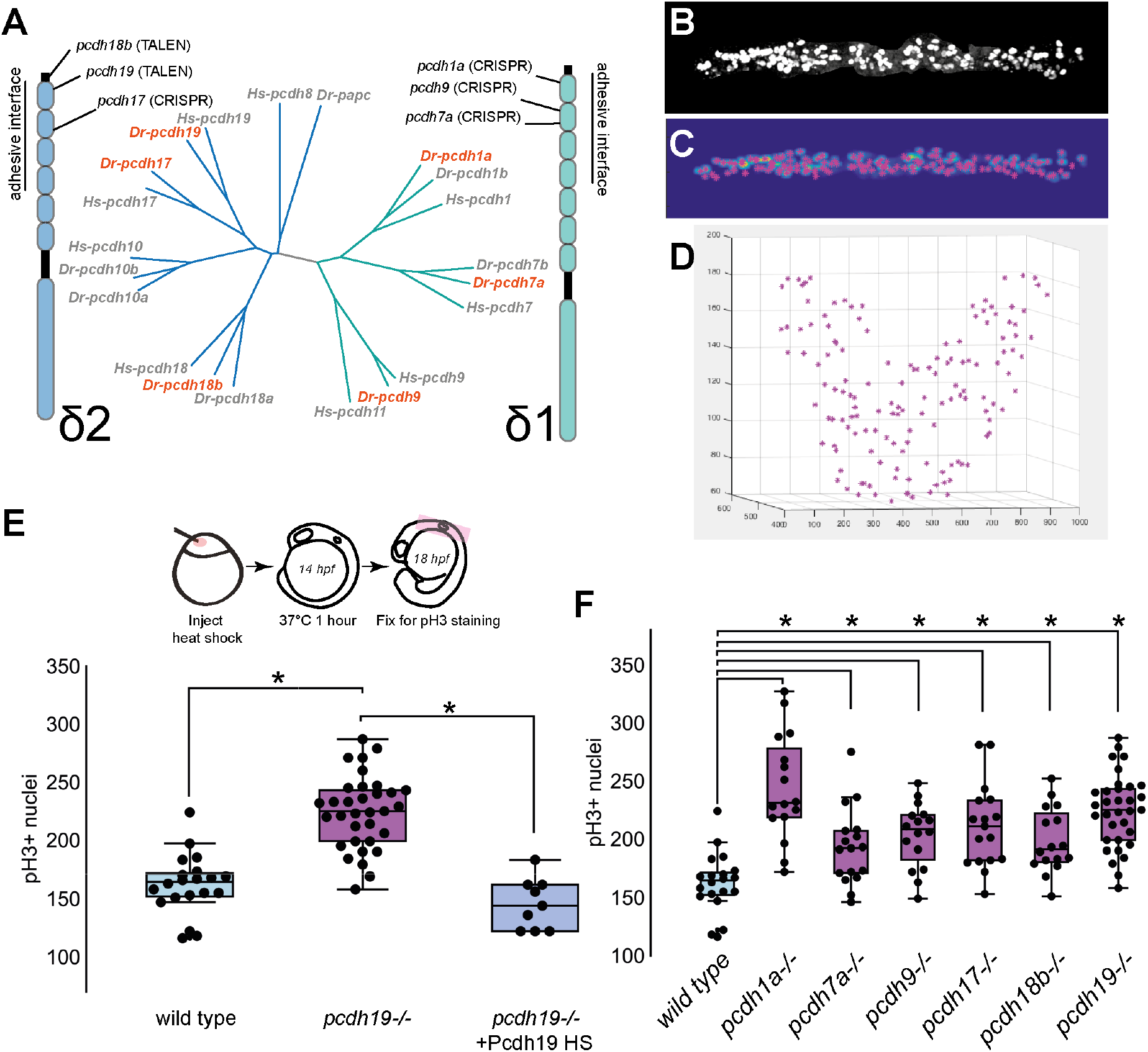
The δ-pcdhs regulate proliferation of neuroepithelial cells. **A**. Overview of the δ-pcdhs. The δ-pcdhs are divided into two subfamilies, the δ1- and δ2-pcdhs, which differ by the presence of 7 or 6 extracellular cadherin repeats, respectively. In both cases, the adhesive interface includes the four distal extracellular domains (as shown). Mutations were generated for six δ-pcdhs (shown in red) and the target sites for each are highlighted. All lesions are within the first two extracellular domains. **B**. A maximum-intensity projection of a wild type 18 hpf embryo labeled with an antibody against phospho-HistoneH3. Analysis was restricted to the neural tube by manually masking out nuclei on the skin or yolk. **C. D**. Labeled cells are identified by 3D cross-correlation. **E**. The number of pH3+ nuclei is elevated in pcdh19 mutants relative to wild type embryos (WT, n=25; pcdh19-/-, n=35; p<0.0001), but is rescued by forced expression of Pcdh19 in the pcdh19 mutants (pcdh19-/-, n=35; pcdh19 hs, n=9; p=0.001). **F**. Each δ-pcdh mutant shows elevated pH3 labeling, relative to wild type embryos. (WT, n=25; pcdh1a-/-, n=24, p<0.0001; pcdh7a-/-, n=21, p=0.0097; pcdh9-/-, n=20, p=0.0003; pcdh17-/-, n=17, p<0.0001; pcdh18b-/-, n=20, p<0.0001; pcdh19-/-, n=35, p<0.0001).

The canonical Wnt/β-catenin signaling pathway plays an important role in regulating cell proliferation and differentiation during development, and mutations in components of Wnt/β-catenin signaling contribute to cancers (). As some studies in cancer cell lines suggest that δ-pcdhs may act as tumor suppressors by regulating β-catenin, we determined the effects of δ-pcdh loss on canonical Wnt/β-catenin signaling. Stimulation of the Wnt/β-catenin signaling pathway results in accumulation of β-catenin, which allows it to translocate to the nucleus and associate with the transcription factor TCF/LEF to activate target genes. We used droplet digital PCR (ddPCR) to quantify the transcript levels of β-catenin/TCF target genes, *axin2* and *lef1* (**Figure 2A,B**). The levels of these transcripts were normalized against the level of *gapdh*. In the case of each δ-pcdh mutant, we observed increased expression (>3-fold) of *axin2* (**Figure 2A**) and *lef1* (**Figure 2B**). Taken together, these data support the idea that canonical Wnt/β-catenin signaling is elevated in the neuroepithelium of zebrafish lacking individual δ-pcdhs.

**Figure 2.**
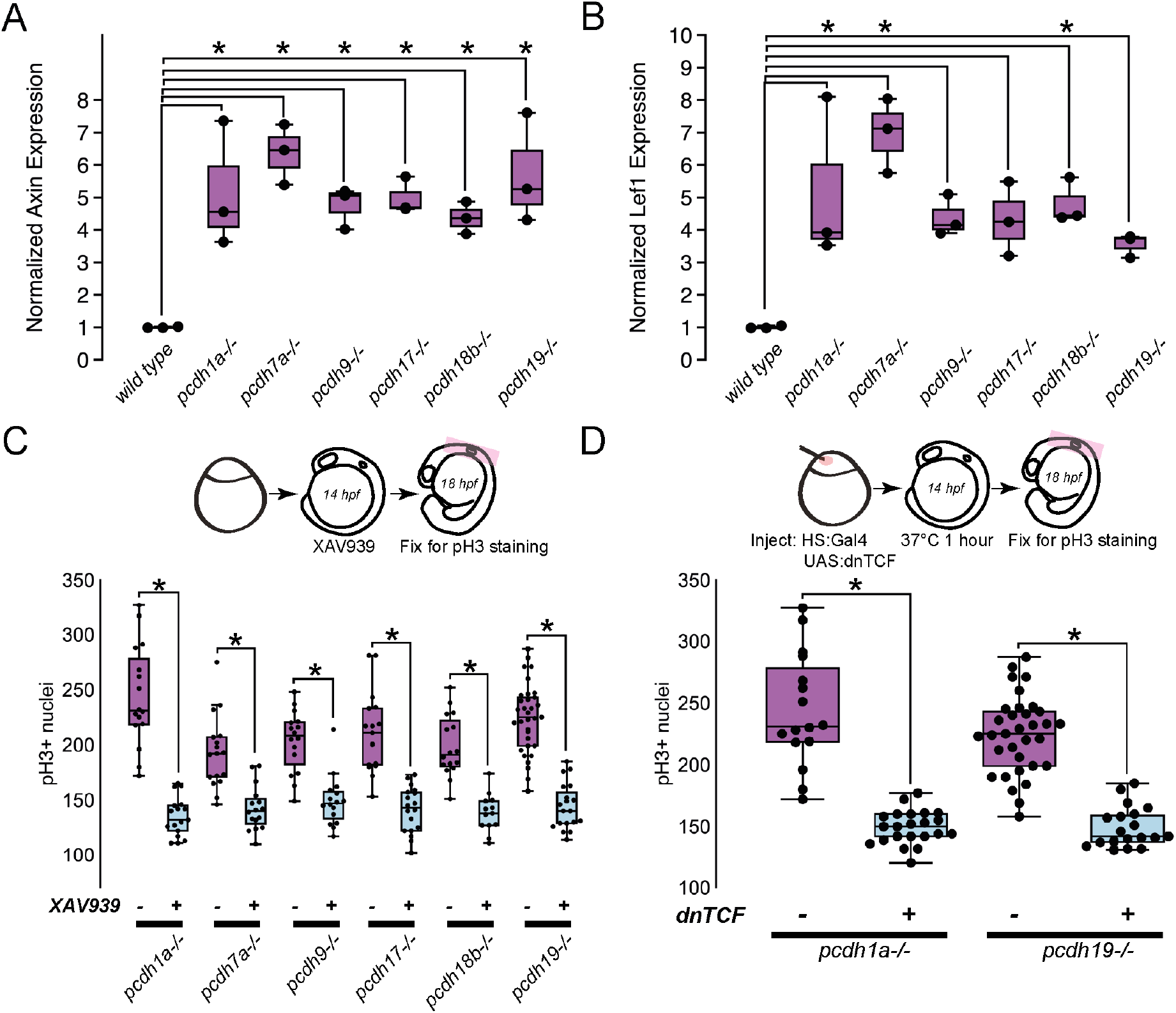
The δ-pcdhs coordinate Wnt/β-catenin signaling to regulate cell proliferation. **A,B**. The expression of Wnt/β-catenin target genes axin2 (**A**) and lef1 (**B**) are elevated >2-fold in each of the δ-pcdh mutants. At least three biological replicates were collected for each genotype. Expression was normalized against gapdh. Expression of axin2 or lef1 in each δ-pcdh line was normalized against the average levels in wild type embryos. (**A**; axin2; WT, n=3; pcdh1a-/-, n=3, p=0.0054; pcdh7a-/-, n=3, p=0.0005; pcdh9-/-, n=3, p=0.0127; pcdh17-/-, n=3, p<0.0079; pcdh18b-/-, n=3, p<0.0278; pcdh19-/-, n=3, p=0.0018). (**B**; lef1; WT, n=3; pcdh1a-/-, n=3, p=0.0128; pcdh7a-/-, n=3, p=0.0006; pcdh9-/-, n=3, p=0.0772; pcdh17-/-, n=3, p<0.0615; pcdh18b-/-, n=3, p<0.0251; pcdh19-/-, n=3, p=0.2175). **C**. To assess the role of Wnt/β-catenin signaling embryos were soaked in 15 μMXAV939, a tankyrase inhibitor that stabilizes axin and inhibits Wnt/β-catenin signaling. In each δ-pcdh mutant, XAV939 eliminates the increased proliferation observed in the mutants. (pcdh1a-/-, n=25; pcdh1a-/- +XAV939, n=17, p<0.0001; pcdh7a-/-, n=21; pcdh7a-/- +XAV939, n=16, p<0.0001; pcdh9-/-, n=20,; pcdh9-/- +XAV939, n=16, p<0.0001; pcdh17-/-, n=17; pcdh17-/- +XAV939, n=18, p<0.0001; pcdh18b-/-, n=20; pcdh18b-/- +XAV939, n=13, p<0.0001; pcdh19-/-, n=35; pcdh19-/- +XAV939, n=19, p<0.0001). **D**. As an alternative approach to blocking Wnt/β-catenin signaling, a dominant-negative TCF (dnTCF) was expressed, in which the amino-terminal β-catenin binding domain was replaced with an HA epitope tag. When expressed in either pcdh1a (a δ1-pcdh) or pcdh19 (a δ2-pcdh) mutants, dnTCF eliminated the increased proliferation that occurs in δ-pcdh mutants. (pcdh1a-/-, n=25; pcdh1a-/- +dnTCF, n=22, p<0.0001; pcdh19-/-, n=35; pcdh19-/- +dnTCF, n=19, p<0.0001

If the increased proliferation observed in δ-pcdh mutants is due to increased activation of canonical Wnt signaling, then blocking Wnt/β-catenin signaling should occlude the proliferation phenotype. To inhibit the canonical Wnt/β-catenin signaling pathway, we treated embryos with the tankyrase inhibitor XAV939 (Huang et al., 2009), which stabilizes Axin, a core component of the β-catenin destruction complex. We treated wild type or δ-pcdh mutant embryos with XAV939 for four hours prior to fixation at 18 hpf and pH3 immunocytochemistry (**Figure 2C**). For each δ-pcdh mutant, treatment with XAV939 blocked the observed increase in cell proliferation (**Figure 2C**). To further probe the involvement of Wnt/β-catenin signaling in the δ-pcdh mutant phenotype, we used a dominant-negative TCF7l1a (dnTCF)(Lewis et al., 2004) to block activation of Wnt target genes (**Figure 2D**). A heat shock driver plasmid *pCS2-hsp70:Gal4-VP16* was co-injected with *pISceI-5xUAS:dnTCF-HA* into wild type and δ-pcdh mutant embryos at the 1-cell stage and embryos were incubated at 37°C from 14-15 hpf, then fixed at 18 hpf. Expression of dnTCF was verified with anti-HA immunocytochemistry. Expression of dnTCF blocked the increased cell proliferation in the hindbrains of δ-pcdh mutants (**Figure 2D**). The results of treatment with XAV939 and expression of dnTCF are consistent with the conclusion that the enhanced proliferation in δ-pcdh mutants is due to elevated Wnt/β-catenin signaling.

Our data support the idea that the δ-pcdhs negatively regulate the canonical Wnt/β-catenin signaling pathway in the zebrafish neuroepithelium. The non-canonical Wnt receptor Ryk is a single-pass transmembrane protein with an extracellular WIF domain and a non-catalytic intracellular kinase domain (Hovens et al., 1992; Lu et al., 2004). Ryk plays important roles in axon guidance (Bonkowsky et al., 1999; Callahan et al., 1995; Li et al., 2009; Schmitt et al., 2006), and has been shown to be required for canonical Wnt/β-catenin signaling in HEK293T cells (Berndt et al., 2011; Lu et al., 2004). Additionally, a proteomics study found several protocadherins to be part of the Ryk interactome (Berndt et al., 2011). To verify the interaction between Ryk and δ-pcdhs, we performed coimmunoprecipitation (co-IP) with extracts of HEK293 cells that had been cotransfected with zebrafish Pcdh18b and zebrafish Ryk (**Figure 3A**). To determine whether this interaction extended to other δ-pcdhs, we cotransfected HEK293 cells with Ryk and δ-pcdhs lacking their intracellular domains (Pcdh1a, Pcdh9, Pcdh17 and Pcdh18b). In each case, the truncated δ-pcdhs co-IPed with Ryk (**Figure 3B**), indicating that the interaction requires the extracellular and/or transmembrane domain of the δ-pcdh. To verify the interaction between Ryk and Pcdh19 *in vivo*, we prepared protein extracts from 18 hpf zebrafish embryos and found that endogenous Pcdh19 was co-IPed with endogenous Ryk (**Figure 3C**). When cotransfected into the ZF4 zebrafish cell line, Pcdh19-GFP and Ryk-HA colocalized both on the cell surface and in intracellular puncta (**Figure 3D**). These data support the idea that δ-pcdh family members interact with Ryk and colocalize in cells, both on the cell surface and in intracellular compartments.

**Figure 3.**
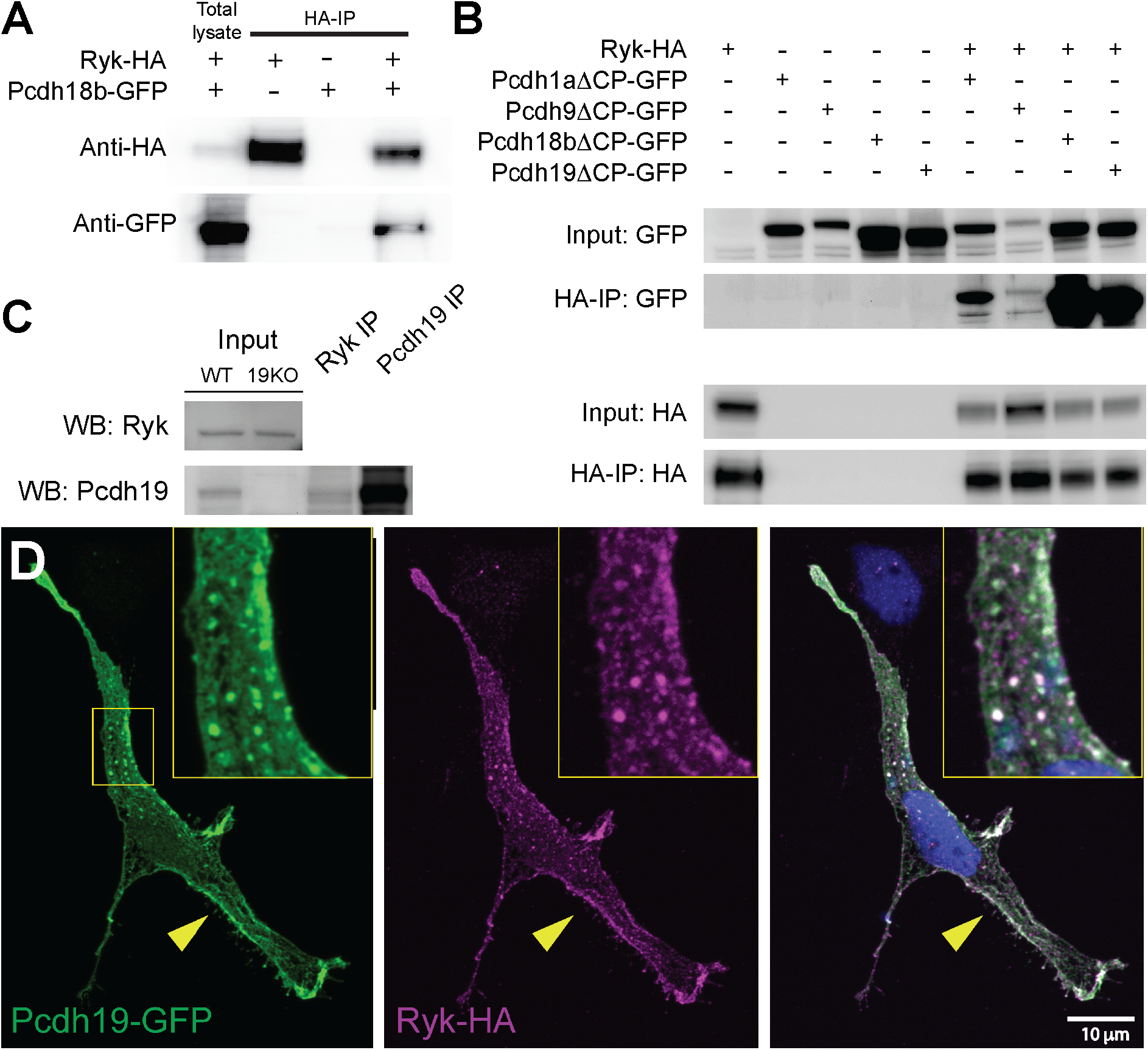
The δ-pcdhs interact with the Wnt receptor Ryk. **A**. When co-transfected into HEK293 cells, zebrafish Ryk-HA is able to co-immunoprecipitate Pcdh18b-GFP. **B**. Ryk-HA is able to co-immunoprecipitate δ1-pcdh (Pcdh1a or Pcdh9) or δ2-pcdh (Pcdh18b or Pcdh19) family members. To improve expression of some δ-pcdhs, we generated constructs lacking most of their intracellular domains (ΔCP). Thus, the intracellular domains are dispensible for interacting with Ryk. **C**. To verify that δ-pcdhs interact with Ryk in vivo, an antibody against endogenous Ryk was used to pull down Ryk from 18 hpf embryo extracts. Western blots show that endogenous Ryk can co-IP endogenous Pcdh19 in vivo.

Based on the close association of the δ-pcdhs with Ryk, both *in vitro* and *in vivo*, we hypothesized that Ryk was required for the enhanced proliferation observed in δ-pcdh mutants. To investigate the role of Ryk in the neuroepithelium, we used CRISPR/Cas9 to generate a mutant zebrafish line lacking *ryk*. Targeting a site near the amino-terminus of zebrafish Ryk, we obtained the frameshift allele, *ryk*(Δ-19) (**Figure 4A**). To determine the effect of Ryk loss on cell proliferation in the neuroepithelium, we performed wholemount immunocytochemistry with antibodies against pH3 in the hindbrain at 18 hpf in *ryk* mutants and in embryos injected with CRISPR/Cas9 directed against *ryk*. In both cases, loss of Ryk resulted in reduced proliferation (**Figure 4B,C**). As prior studies have suggested that Ryk is required for canonical Wnt/β-catenin signaling *in vitro*, we used ddPCR to determine the effects of Ryk loss on the levels of the β-catenin/TCF target genes, *axin2* and *lef1* (**Figure 4D,E**). We found that the Wnt/β-catenin pathway was suppressed in *ryk* mutant embryos and in embryos injected with CRISPR/Cas9 directed against *ryk* (**Figure 4D,E**). This is consistent with Ryk participating in canonical Wnt signaling in the developing zebrafish neuroepithelium.

**Figure 4.**
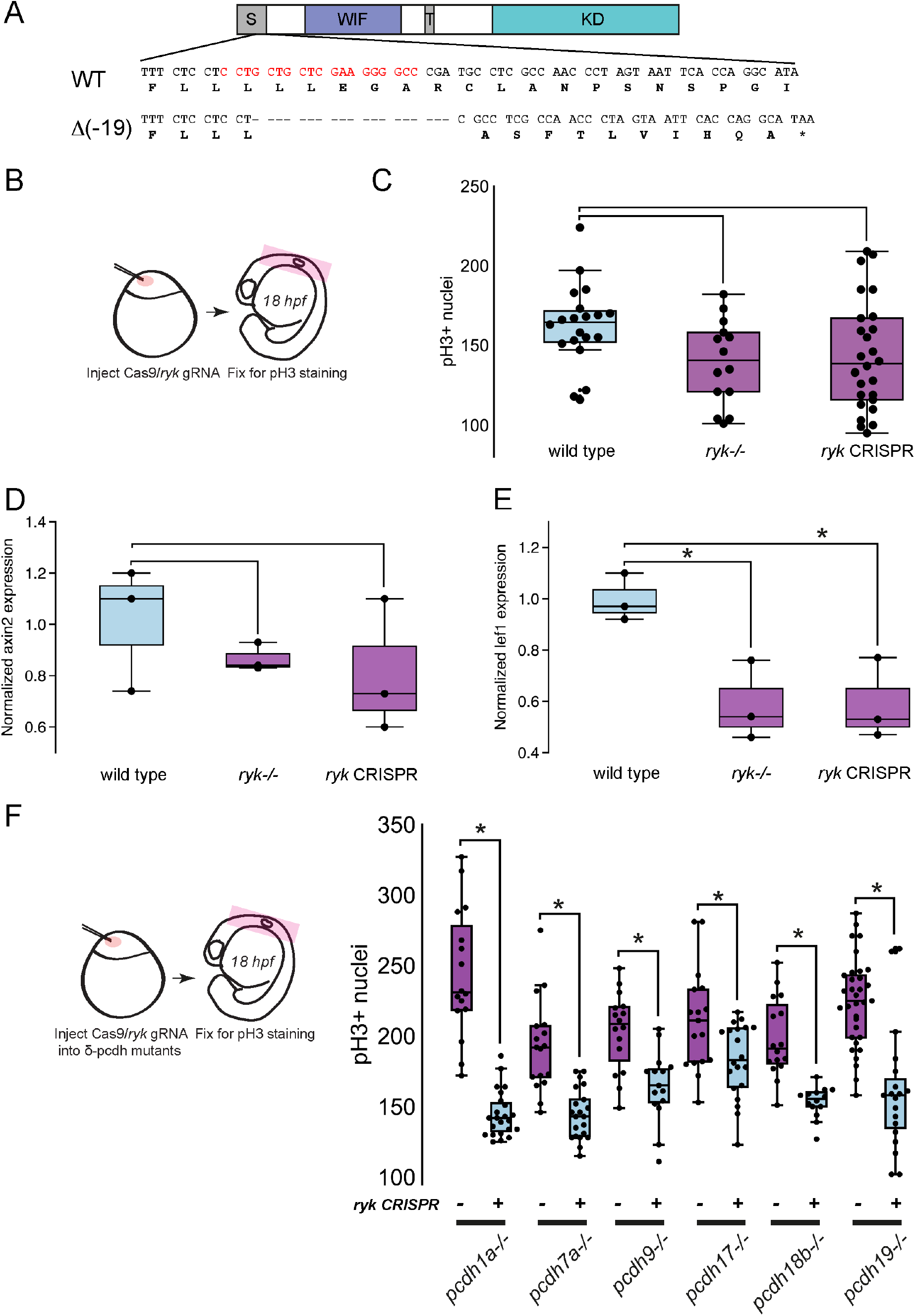
Ryk promotes Wnt/β-catenin signaling and is required for increased cell proliferation in δ-pcdh mutants. **A**. CRISPR/Cas9 was used to generate indel mutations in zebrafish ryk. The selected target site (red text) was within the sequence encoding the signal peptide. Two frameshift mutations were identified that result in a premature stop codon within the distal extracellular domain. **B,C**. Loss of ryk shows a modest decrease in pH3 labeling in 18 hpf zebrafish embryos. In addition to being used for generating germline lesions in ryk, the gRNA directed against ryk is a very effective knockdown reagent. Both ryk mutants and ryk CRISPR injected embryos showed a modest decrease in pH3 labeling, which was not statistically significant. **D,E**. Loss or reduction of ryk leads to reduced levels of the Wnt/β-catenin target genes axin2 (**D**) and lef1 (**E**) as determined by ddPCR, suggesting that Ryk promotes canonical Wnt signaling. (lef1: wt vs ryk-/-,n=3, p=0.03; lef1: wt-ryk CRISPR,n=3, p=0.0275). **F**. CRISPR/Cas9 was used to knockdown Ryk expression in each of the δ-pcdh mutant lines. Reduction of Ryk in the mutant backgrounds eliminated the increased pH3 labeling, suggesting that the presence of Ryk is required for the increased proliferation in these lines. (pcdh1a-/-, n=25; pcdh1a-/- +rykCRISPR, n=17, p<0.0001; pcdh7a-/-, n=23; pcdh7a-/- + rykCRISPR, n=20, p<0.0001; pcdh9-/-, n=20,; pcdh9-/- + rykCRISPR, n=15, p<0.0001; pcdh17- /-, n=17; pcdh17-/- + rykCRISPR, n=21, p<0.0001; pcdh18b-/-, n=20; pcdh18b-/- + rykCRISPR, n=14, p<0.0001; pcdh19-/-, n=35; pcdh19-/- + rykCRISPR, n=19, p<0.0001).

The loss of δ-pcdhs leads to enhanced Wnt/β-catenin signaling and cell proliferation (**Figure 2**). In contrast, the loss of Ryk results in reduced Wnt/β-catenin signaling and cell proliferation (**Figure 4D,E**). As we show that Ryk can exist in a complex with δ-pcdhs (**Figure 3**), we hypothesized that Ryk might be required for the δ-pcdh phenotype. To test this, we used CRISPR/Cas9 injection to knockdown Ryk in wild type and δ-pcdh mutant embryos (**Figure 4F**). In each case, knockdown of Ryk blocked the increased proliferation observed in δ-pcdh mutants (**Figure 4F**). These results suggest that the presence of Ryk is required for the enhanced proliferation resulting from the loss of δ-pcdhs. As Ryk participates in the canonical Wnt/β-catenin pathway, we speculate that: 1) δ-pcdhs negatively regulate Ryk, possibly by regulating its trafficking or stability, and 2) the increased Wnt/β-catenin signaling observed in δ-pcdh mutants is due to disinhibition of Ryk.

A previous study showed that zebrafish *pcdh19* is expressed in a subset of neural progenitor cells within the optic tectum, and that cell divisions of these progenitors give rise to radial columns of sibling neurons that share the expression of *pcdh19* (Cooper et al., 2015). Other evidence supports the observation that δ-pcdhs are expressed in neural progenitor cells (Bergsland et al., 2011; McAninch and Thomas, 2014) and that they regulate neurogenesis (Fujitani et al., 2016; Homan et al., 2018; Zhang et al., 2014). Here, we show that the loss of individual δ-pcdh family members leads to increased cell proliferation, and that they likely do so through an increase in Wnt/β-catenin signaling. As δ-pcdhs exhibit distinct regional expression in the nervous system, and may define distinct cell lineages within a brain region, the δ-pcdhs could provide a mechanism to differentially modulate cell responsiveness to canonical Wnt signaling within populations of progenitor cells. Our data support the idea that the δ-pcdhs interact with canonical Wnt signaling through the receptor Ryk, as Ryk is required for the elevated pH3 labeling observed in δ-pcdh mutants. These results are consistent with δ-pcdhs acting as novel upstream regulators of canonical Wnt signaling, and provide insight into the contributions of both δ-pcdhs (Hirano and Takeichi, 2012; Redies et al., 2012) and Wnt/β-catenin signaling (De Ferrari and Moon, 2006; Kwan et al., 2016) to neurodevelopmental disorders and to cancer (Berx and van Roy, 2009; Polakis, 2012; van Roy, 2014; Zhan et al., 2017). Further work will be required to determine the mechanisms of Ryk regulation by δ-pcdhs and Ryk involvement in Wnt signaling, as well as the precise role of differential adhesion by δ-pcdhs.

## Methods

### Fish Maintenance

Adult zebrafish (Danio rerio) and embryos of the Tübingen longfin and AB strains were maintained at ~28.5°C in E3 buffer (5 mM NaCl, 0.17 mM KCl, 0.33 mM CaCl_2_, 0.33 mM MgSO_4_, pH 7.2) and staged according to ((Westerfield, 1995). All zebrafish experiments and procedures were performed in compliance with institutional ethical regulations for animal research at Ohio State University and were approved by the university’s Institutional Animal Care and Use Committee.

### TALEN and CRISPR construction and production of germline lesions

We used the online tool TAL Effector Nucleotide Targeter 2.0 (https://tale-nt.cac.cornell.edu/node/add/talen) to search for a TALEN target site in exon 1 of zebrafish *pcdh18b*. We identified an appropriate target site downstream of the signal peptide, CTGAAACTTATGCTTCTGGCggccgtggcgcacaaTGTTTCGGGGAAGACTTTAA (uppercase indicates TAL left and right binding sites). The TALEN arrays (left: HD-NG-NH-NI-NI-NI-HD-NG-NG-NI-NG-NH-HD-NG-NG-HD-NG-NH-NH-HD and right: NG-NG-NI-NI-NI-NH-NG-HD-NG-NG-HD-HD-HD-HD-NH-NI-NI-NI-HD-NI) were assembled in RCIscript-GoldyTALEN (Bedell et al., 2012) using the TAL Effector Kit 1.0 (Cermak et al., 2011). The plasmid kit used for the generation of TALENs was a gift of Daniel Voytas and Adam Bogdanove (Addgene kit #1000000016). Plasmid encoding assembled TALENs was linearized with SacI, and used as template for mRNA synthesis with a T3 mMessage Machine kit (Ambion). To generate germline lesions in zebrafish *pcdh18b*, we injected 1 cell stage embryos with 50 pg mRNA encoding the left and right nucleases. Injected embryos were grown to adulthood and screened for germline lesions. For screening, adult F0 fish were outcrossed with wild types, and genomic DNA was prepared from eight embryos of each cross. High resolution melt analysis (HRMA) was used to identify putative founders (Dahlem et al., 2012). PCR products exhibiting aberrant melting curves were cloned and sequenced. F0 adults exhibiting frameshift mutations were outcrossed and the F1 embryos were grown to adulthood. To screen the adult F1 fish, genomic DNA was prepared from caudal fin clips screened by HRMA. These heterozygote F1 founders were outcrossed again and the F2 offspring were raised and screened to establish mutant lines. To obtain homozygous *pcdh18b* mutants heterozygotes for each allele were incrossed and embryos were grown to adulthood and screened by HRMA.

Germline mutations for *pcdh1a, pcdh7a, pcdh9* and *pcdh17* were generated using CRISPR/Cas9. Target sites were identified using either ZiFiT (partners.zifit.org/ZiFiT/) or with CRISPRscan (https://www.crisprscan.org). For each target site, top and bottom oligonucleotides were annealed to generate double stranded oligos with overhangs compatible with the BsaI sites of plasmid DR274 (Hwang et al., 2013). DR274 was a gift from Keith Joung (Addgene plasmid #42250; http://n2t.net/addgene:42250; RRID:Addgene_42250). Plasmids encoding target site gRNAs were verified by Sanger sequencing. Each gRNA plasmid was linearized with DraI and used as template for RNA synthesis with a T7 MAXIscript kit (Ambion). Plasmid pCS2-nCas9n (Jao et al., 2013) encoding Cas9 was linearized with NotI and used as template for mRNA synthesis using a SP6 mMessage Machine kit (Ambion). pCS2-nCas9n was a gift from Wenbiao Chen (Addgene plasmid #47929; http://n2t.net/addgene:47929; RRID:Addgene_47929). One-cell stage embryos were injected with 1 nL of 80ng/μL of Cas9 mRNA and 40 ng/μL gRNA. Injected embryos were grown to adulthood and screened for germline lesions, as described above.

### Whole mount in situ hybridization

Riboprobes directed against δ-protocadherin extracellular domains were synthesized by first amplifying approximately 1 Kb of the extracellular domain by PCR using 3 dpf cDNA, as described previously (Biswas et al., 2014b; Biswas and Jontes, 2009; Blevins et al., 2011; Emond et al., 2009). A T7 RNA Polymerase binding site was included in each of the reverse primers, and these PCR products were used as templates for *in vitro* transcription (Promega). Antisense riboprobes were labeled with digoxygenin-dUTP or fluorescein-dUTP (Roche). Whole mount *in situ* hybridizations were carried out using standard methods (Westerfield, 1995). Briefly, embryos were fixed at 4°C overnight in 4% paraformaldehyde in PBS, dehydrated in a methanol series and stored in 100% methanol overnight at −20°C. They were rehydrated in decreasing concentrations of methanol and embryos 24 hpf and older were permeabilized using Proteinase K (10μg/ml, Roche). Embryos were refixed in 4% paraformaldehyde prior to hybridization. Labeled riboprobe was added to the hybridization buffer at a final concentration of 200 ng/ml and hybridization was carried out at 65°C overnight. Alkaline phosphatase-conjugated anti-digoxygenin Fab fragments (Roche) were used at 1:5000 dilution. NBT/BCIP (Roche) was used for the coloration reaction. Images were captured on a Leica MZ16F stereomicroscope (Leica Microsystems). For double fluorescent *in situ* hybridization, embryos were fixed and hybridized with riboprobes against δ-protocadherins (labeled with digoxygenin-dUTP) and *her4.1* (labeled with fluorescein-dUTP). The digoxygenin-labeled probe was detected using anti-digoxygenin Fab-POD (Roche), and developed using the tetramethylrhodamine substrate from the TSA Plus kit (Perkin-Elmer). Subsequently, the fluorescein probe was detected with an anti-fluorescein primary antibody (Roche) and a goat anti-mouse-HRP secondary antibody (Invitrogen), and developed using the fluorescein substrate from the TSA Plus kit. Embryos were imaged using two-photon microscopy as described above. Maximum projection images of 50 optical sections from each channel were generated in FIJI using the Smooth Manifold Extraction plug-in (Shihavuddin et al., 2017).

### Coimmunoprecipitation and Western blotting

HEK293 cells (CRL-1573, American Type Culture Collection) were transiently transfected with plasmids encoding GFP- or HA-tagged zebrafish protocadherins and Ryk using calcium phosphate precipitation as described previously (Cooper et al., 2015). After 24 hours, cells were rinsed in PBS and lysed on ice in cell lysis buffer (CLB) (20 mM Tris, pH 7.5, 150 mM NaCl, 1 mM EDTA, 0.5% Triton X-100, 1 mM PMSF, and 1X cOmplete protease inhibitor cocktail (Roche) and microcentrifuged at 4°C for 10 min. Supernatants were incubated with anti-HA magnetic beads (Thermo Fisher Scientific) overnight at 4°C. For GFP coimmunoprecipitation, the lysates were incubated for 1 hour with 2ug anti-GFP antibody (Thermo Fisher Scientific) and then incubated for 30-60 minutes with Protein A Dynabeads (Thermo Fisher Scientific). The beads were washed five times in wash buffer (20 mM Tris, pH 7.5, 150 mM NaCl, 0.5% Triton X-100), resuspended in loading buffer, and heated to 70°C for 10 minutes. Samples were loaded onto 10% Bis-Tris NuPAGE gels (Thermo Fisher Scientific) and subjected to electrophoresis. Proteins were then transferred (Bio-Rad Laboratories) to PVDF (GE Life Science), blocked with 5% nonfat milk in TBST, and incubated overnight with primary antibody (Invitrogen rabbit anti-GFP, 1:1,000; Invitrogen mouse anti-HA, 1:5,000). HRP-conjugated secondary antibodies (Jackson ImmunoResearch Laboratories) were used at 1:5,000, and the chemiluminescent signal was amplified using Western Lightning Ultra (PerkinElmer). Blots were imaged on a molecular imaging system (Omega 12iC; UltraLum, Inc.). For the *in vivo* coimmunoprecipitation experiments, 18 hpf embryos were lysed in CLB, as described above, and incubated with either 2 μg rabbit anti-Ryk (Proteintech) or rabbit anti-Pcdh19 (custom made by Covance; (Biswas et al., 2010). Proteins were coimmunoprecipitated using Protein A Dynabeads (Thermo Fisher Scientific) and Western blots were performed as described above (1:1000 used for both anti-Ryk and anti-Pcdh19 primary antibodies for Western blots).

### ZF4 cell culture and immunocytochemistry

The zebrafish fibroblast cell line ZF4 (CRL-2050, American Type Culture Collection) was cultured in DMEM: F12 media (GIBCO) with 10% fetal bovine serum and maintained at 28°C. ZF4 cells were seeded on glass coverslips and transfected with plasmids encoding Pcdh19-GFP and Ryk-HA using Fugene HD (Promega) according to the manufacturer’s protocol. Cells were fixed 24h later with 4% paraformaldehyde in PBS for 10 min, permeabilized with 0.25% Triton X-100/PBS for 5 min, and rinsed thoroughly. Cells were then incubated overnight at 4°C in blocking solution (PBS, 2% normal goat serum, 3% BSA) and primary antibodies (mouse anti-HA, Invitrogen; rabbit anti-GFP, Invitrogen). Cells were then immunolabeled with Alexa-conjugated secondary antibodies (Thermo Fisher Scientific) and DAPI (Thermo Fisher Scientific) was added to visualize cell nuclei. Coverslips were rinsed and mounted in Fluoromount G (Electron Microscopy Science). Images were captured on an Andor spinning disk confocal system fitted with a Nikon TiE inverted epifluorescence microscope equipped with a 100x/1.4NA Plan-Apochromat VC oil immersion objective, a Lumencor SOLA LED light source for epifluorescence illumination, and an Andor iXon Ultra 897 EMCCD camera.

### Axin2 and Lef1 transcript quantification

Total RNA was extracted from 18 hpf embryos using TRI Reagent (Sigma-Aldrich) and cDNA was synthesized using SuperScript IV First-Strand Synthesis System (Thermo Fisher Scientific). Digital droplet PCR (ddPCR) was used to quantify Axin2 and Lef1 transcript levels in mutant and WT embryos and normalized to GAPDH expression. In brief, 15000-18000 droplets containing cDNA, primers for either Axin2 or Lef1 and GAPDH, probes for either Axin2 or Lef1 and GAPDH, 2X ddPCR SuperMix (Bio-Rad) and droplet generating oil were generated with QX200 droplet generator (Bio-Rad).

Primers (Eurofins genomics):

Axin2-F 5’ CGGACACTTCAAGGAACAACTACG 3’
Axin2-R 5’ TGCCCTCATACATTGGCAGAACTG 3’
Lef1-F 5’ GAGGGAAAAGATCCAGGAAC 3’
Lef1-R 5’ AGGTTGAGAAGTCTAGCAGG 3’
GAPDH-F 5’ AAGTGTCAGGACGAACAGAG 3’
GAPDH-R 5’ GCGACCGAATCCGTTAATAC 3’

Probes (Eurofins genomics):

Axin FAM-TGATGAGTTTGAGTGTGGTGCTGTGTTCGA-BHQ1
Lef1 FAM-AACAATGGAGGAAAGCGGACGTCTT-BHQ1
GAPDH HEX–ACAAACGAGGACACAACCAAATCAGG-BHQ1

cDNA partitioned within these droplets were PCR amplified and fluorescence from these droplets detected by the QX200 droplet reader (Bio-Rad). The relative abundance of Axin2 and Lef1 transcripts was calculated using Poisson statistical distribution of fluorescence absorption and normalized to GAPDH transcript level using QuantaSoft software (Bio-Rad). Two technical replicates and at least three biological replicates were performed for each sample.

### Immunocytochemistry and two-photon imaging

Zebrafish embryos were fixed for 1 hour in 4% paraformaldehyde in PBS, then rinsed thoroughly in PBS before permeabilizing in acetone at −20°C for 7 minutes. Embryos were then rinsed in water and then PBS + 0.1% Tween-20 (PBST). Embryos were then blocked in PBST, 2% Roche blocking reagent, 1% DMSO, and 10% fetal bovine serum. Primary antibodies were added to the samples for overnight incubation (rabbit anti-phospho-Histone H3 (1:200; Cell Signaling); mouse anti-HA (1:500; Invitrogen)). Alexa 488 and 594 secondary antibodies were used at 1:500 (Thermo Fisher Scientific). Two-photon imaging of embryos was performed at room temperature on a custom-built resonant-scanning microscope (Light and Jontes, 2019) controlled by ScanImage (Pologruto et al., 2003). Excitation was provided by a Chameleon-XR Ti:Sapphire laser (Coherent, Inc.) tuned to 800nm. We used a Nikon 25x/1.1NA water immersion objective. Embryos were oriented in low-melting point agarose (Sigma Aldrich) and immersed in PBS. Image stacks of 128 optical sections were taken at 1 μm intervals (1024 x 1024 pixels). To assess the role of Wnt/β-catenin signaling in cell proliferation, embryos were treated with the inhibitor XAV939 (Cayman Chemical). The inhibitor was added to the embryos at a final concentration of 15 μM in E3 buffer starting at 14 hpf prior to fixation at 18 hpf for phospho-Histone H3 immunolabeling. In a subset of experiments, embryos were injected at the 1-cell stage with 25 ng/μL pCS2:hsp:Gal4-VP16 and 25 ng/μL pISceI:5XUAS:HA-dnTCF or pISceI:5XUAS:Pcdh19-GFP plasmids. The expression of HA-dnTCF or Pcdh19-GFP was initiated by heat shocking the embryos at 37°C for 1 hour at 14 hpf, and the embryos were allowed to recover before fixation at 18 hpf for phospho-Histone H3 immunocytochemistry. In the case of HA-dnTCF, the embryos were sorted for strong anti-HA fluorescence prior to mounting and imaging.

### Image analysis and cell counting

Images stacks were initially adjusted for brightness and contrast in Fiji (Schindelin et al., 2012). Maximum intensity projections were made to determine the boundary of the neural tube and a manual mask was drawn to exclude non-neural nuclei from the analysis. For cross-correlation, a template nucleus was generated by cropping a 16×16×32 region around a well-labeled, well-isolated nucleus from one of the image stacks. Automated cell counting was performed in MATLAB (www.mathworks.com). Both the template and the image stacks were padded to 1024×1024×256 and a 3D cross-correlation was performed. An initial threshold for the correlation peaks was set manually to detect as many nuclei as possible and to eliminate any spurious peaks. Once set, this threshold was used for all analyzed image stacks.

### Statistical Analysis

For pairwise comparisons, a two-tailed Student’s t-test was used. For comparisons among groups, a one-way ANOVA with Tukey HSD was used. Statistics were calculated either with JMP Pro 14 (www.jmp.com) or with GraphPad Prism 6 (www.graphpad.com).

## Acknowledgements

This work was supported by the National Science Foundation Grant IOS 1457126, an NIH Grant R01 EY027003, NIH Shared Instrumentation Grants S10 OD010383, S10 OD18056, P30 CA016058, and an NIH Neurosciences Center Core Grant P30 NS104177. We thank Harold Ortiz-Medina and Min An for technical assistance in generating mutant lines. We thank K. Joung, W.Chen, D. Voytas and A. Bogdanove for reagents, and V. McGovern for help with ddPCR.

---

**Supplemental Figure 1.**
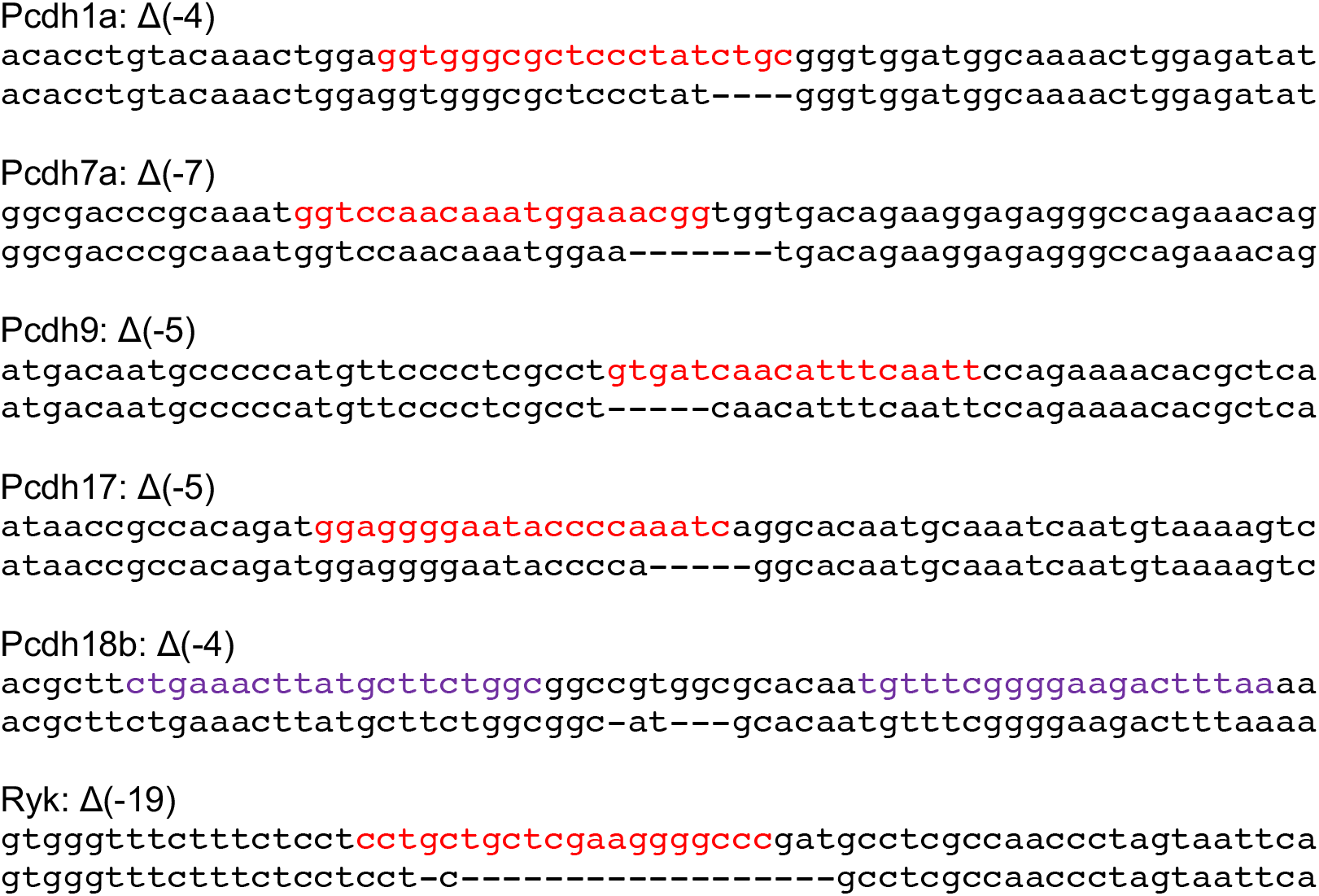
Germline target sites and lesions. Shown are alignments of genomic sequences for wild type and mutant embryos, for each of the mutant lines generated in this study: pcdh1a, pcdh7a, pcdh9, pcdh17 and pcdh18b. CRISPR target sites are shown in red. The TALEN binding sites for pcdh18b are shown in purple.

**Supplemental Figure 2.**
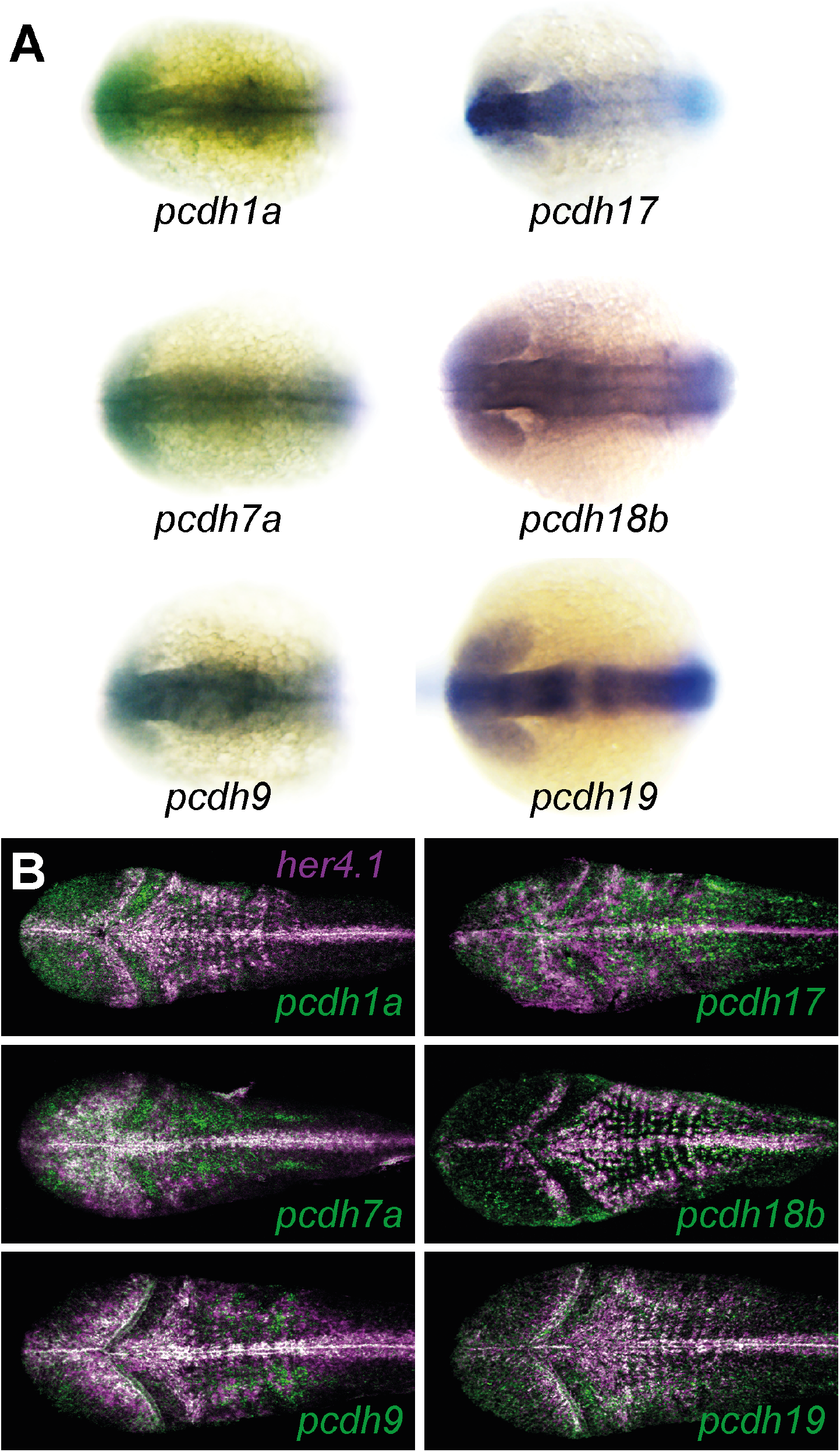
The δ-pcdhs are expressed in the zebrafish neuroepithelium. **A**. in situ hybridization using riboprobes directed against pcdh1a, pcdh7a, pcdh9, pcdh17, pcdh18b and pcdh19. Each of these δ-pcdhs is expressed in the neuroepithelium of 18 hpf zebrafish embryos. **B**. At later stages, δ-pcdhs are expressed both in neural progenitor cells and in neurons. Shown here are maximum intensity projection images of the optic tecta and hindbrains of 36 hpf embryos labeled with riboprobes against her4.1 (magenta) and δ-pcdhs (green). Overlap (white) is greatest along the midline, the posterior wall of the optic tectum.

## REFERENCES

Aamar, E., and I.B. Dawid. 2008. Protocadherin-18a has a role in cell adhesion, behavior and migration in zebrafish development. Dev Biol. 318:335–346.

Bedell, V.M., Y. Wang, J.M. Campbell, T.L. Poshusta, C.G. Starker, R.G. Krug, 2nd, W. Tan, S.G. Penheiter, A.C. Ma, A.Y. Leung, S.C. Fahrenkrug, D.F. Carlson, D.F. Voytas, K.J. Clark, J.J. Essner, and S.C. Ekker. 2012. In vivo genome editing using a high-efficiency TALEN system. Nature. 491:114–118.

Bergsland, M., D. Ramskold, C. Zaouter, S. Klum, R. Sandberg, and J. Muhr. 2011. Sequentially acting Sox transcription factors in neural lineage development. Genes Dev. 25:2453–2464.

Berndt, J.D., A. Aoyagi, P. Yang, J.N. Anastas, L. Tang, and R.T. Moon. 2011. Mindbomb 1, an E3 ubiquitin ligase, forms a complex with RYK to activate Wnt/beta-catenin signaling. J Cell Biol. 194:737–750.

Berx, G., and F. van Roy. 2009. Involvement of members of the cadherin superfamily in cancer. Cold Spring Harb Perspect Biol. 1:a003129.

Bing, Y., M. Tian, G. Li, B. Jiang, Z. Ma, L. Li, L. Wang, H. Wang, and D. Xiu. 2018. Down-regulated of PCDH10 predicts poor prognosis in hepatocellular carcinoma patients. Medicine (Baltimore). 97:e12055.

Bisogni, A.J., S. Ghazanfar, E.O. Williams, H.M. Marsh, J.Y. Yang, and D.M. Lin. 2018. Tuning of delta-protocadherin adhesion through combinatorial diversity. Elife. 7.

Biswas, S., M.R. Emond, P.Q. Duy, T. Hao le, C.E. Beattie, and J.D. Jontes. 2014a. Protocadherin-18b interacts with Nap1 to control motor axon growth and arborization in zebrafish. Mol Biol Cell. 25:633–642.

Biswas, S., M.R. Emond, P.Q. Duy, L.T. Hao, C.E. Beattie, and J.D. Jontes. 2014b. Protocadherin-18b interacts with Nap1 to control motor axon growth and arborization in zebrafish. Mol Biol Cell. 25:633–642.

Biswas, S., M.R. Emond, and J.D. Jontes. 2010. Protocadherin-19 and N-cadherin interact to control cell movements during anterior neurulation. J Cell Biol. 191:1029–1041.

Biswas, S., and J.D. Jontes. 2009. Cloning and characterization of zebrafish protocadherin–17. Dev Genes Evol. 219:265–271.

Blevins, C.J., M.R. Emond, S. Biswas, and J.D. Jontes. 2011. Differential expression, alternative splicing, and adhesive properties of the zebrafish delta1-protocadherins. Neuroscience. 199:523–534.

Bonkowsky, J.L., S. Yoshikawa, D.D. O’Keefe, A.L. Scully, and J.B. Thomas. 1999. Axon routing across the midline controlled by the Drosophila Derailed receptor. Nature. 402:540–544.

Bruining, H., A. Matsui, A. Oguro-Ando, R.S. Kahn, H.M. Van’t Spijker, G. Akkermans, O. Stiedl, H. van Engeland, B. Koopmans, H.A. van Lith, H. Oppelaar, L. Tieland, L.J. Nonkes, T. Yagi, R. Kaneko, J.P. Burbach, N. Yamamoto, and M.J. Kas. 2015. Genetic Mapping in Mice Reveals the Involvement of Pcdh9 in Long-Term Social and Object Recognition and Sensorimotor Development. Biol Psychiatry. 78:485–495.

Cadigan, K.M., and Y.I. Liu. 2006. Wnt signaling: complexity at the surface. J Cell Sci. 119:395–402.

Callahan, C.A., M.G. Muralidhar, S.E. Lundgren, A.L. Scully, and J.B. Thomas. 1995. Control of neuronal pathway selection by a Drosophila receptor protein-tyrosine kinase family member. Nature. 376:171–174.

Cermak, T., E.L. Doyle, M. Christian, L. Wang, Y. Zhang, C. Schmidt, J.A. Baller, N.V. Somia, A.J. Bogdanove, and D.F. Voytas. 2011. Efficient design and assembly of custom TALEN and other TAL effector-based constructs for DNA targeting. Nucleic Acids Res. 39:e82.

Chen, X., and B.M. Gumbiner. 2006. Paraxial protocadherin mediates cell sorting and tissue morphogenesis by regulating C-cadherin adhesion activity. J Cell Biol. 174:301–313.

Clevers, H. 2006. Wnt/beta-catenin signaling in development and disease. Cell. 127:469–480.

Cooper, S.R., M.R. Emond, P.Q. Duy, B.G. Liebau, M.A. Wolman, and J.D. Jontes. 2015. Protocadherins control the modular assembly of neuronal columns in the zebrafish optic tectum. J Cell Biol. 211:807–814.

Cooper, S.R., J.D. Jontes, and M. Sotomayor. 2016. Structural determinants of adhesion by Protocadherin-19 and implications for its role in epilepsy. Elife. 5.

Dahlem, T.J., K. Hoshijima, M.J. Jurynec, D. Gunther, C.G. Starker, A.S. Locke, A.M. Weis, D.F. Voytas, and D.J. Grunwald. 2012. Simple methods for generating and detecting locus-specific mutations induced with TALENs in the zebrafish genome. PLoS Genet. 8:e1002861.

De Ferrari, G.V., and R.T. Moon. 2006. The ups and downs of Wnt signaling in prevalent neurological disorders. Oncogene. 25:7545–7553.

Depienne, C., D. Bouteiller, B. Keren, E. Cheuret, K. Poirier, O. Trouillard, B. Benyahia, C. Quelin, W. Carpentier, S. Julia, A. Afenjar, A. Gautier, F. Rivier, S. Meyer, P. Berquin, M. Helias, I. Py, S. Rivera, N. Bahi-Buisson, I. Gourfinkel-An, C. Cazeneuve, M. Ruberg, A. Brice, R. Nabbout, and E. Leguern. 2009. Sporadic infantile epileptic encephalopathy caused by mutations in PCDH19 resembles Dravet syndrome but mainly affects females. PLoS Genet. 5:e1000381.

Depienne, C., and E. LeGuern. 2012. PCDH19-related infantile epileptic encephalopathy: an unusual X-linked inheritance disorder. Hum Mutat. 33:627–634.

Dibbens, L.M., P.S. Tarpey, K. Hynes, M.A. Bayly, I.E. Scheffer, R. Smith, J. Bomar, E. Sutton, L. Vandeleur, C. Shoubridge, S. Edkins, S.J. Turner, C. Stevens, S. O’Meara, C. Tofts, S. Barthorpe, G. Buck, J. Cole, K. Halliday, D. Jones, R. Lee, M. Madison, T. Mironenko, J. Varian, S. West, S. Widaa, P. Wray, J. Teague, E. Dicks, A. Butler, A. Menzies, A. Jenkinson, R. Shepherd, J.F. Gusella, Z. Afawi, A. Mazarib, M.Y. Neufeld, S. Kivity, D. Lev, T. Lerman-Sagie, A.D. Korczyn, C.P. Derry, G.R. Sutherland, K. Friend, M. Shaw, M. Corbett, H.G. Kim, D.H. Geschwind, P. Thomas, E. Haan, S. Ryan, S. McKee, S.F. Berkovic, P.A. Futreal, M.R. Stratton, J.C. Mulley, and J. Gecz. 2008. X-linked protocadherin 19 mutations cause female-limited epilepsy and cognitive impairment. Nat Genet. 40:776–781.

Emond, M.R., S. Biswas, and J.D. Jontes. 2009. Protocadherin-19 is essential for early steps in brain morphogenesis. Dev Biol. 334:72–83.

Fujitani, M., S. Zhang, R. Fujiki, Y. Fujihara, and T. Yamashita. 2016. A chromosome 16p13.11 microduplication causes hyperactivity through dysregulation of miR-484/protocadherin-19 signaling. Mol Psychiatry.

Harrison, O.J., J. Brasch, P.S. Katsamba, G. Ahlsen, A.J. Noble, H. Dan, R.V. Sampogna, C.S. Potter, B. Carragher, B. Honig, and L. Shapiro. 2020. Family-wide Structural and Biophysical Analysis of Binding Interactions among Non-clustered delta-Protocadherins. Cell Rep. 30:2655–2671 e2657.

Hayashi, S., Y. Inoue, H. Kiyonari, T. Abe, K. Misaki, H. Moriguchi, Y. Tanaka, and M. Takeichi. 2014. Protocadherin-17 mediates collective axon extension by recruiting actin regulator complexes to interaxonal contacts. Dev Cell. 30:673–687.

Hirano, S., and M. Takeichi. 2012. Cadherins in brain morphogenesis and wiring. Physiol Rev. 92:597–634.

Homan, C.C., S. Pederson, T.H. To, C. Tan, S. Piltz, M.A. Corbett, E. Wolvetang, P.Q. Thomas, L.A. Jolly, and J. Gecz. 2018. PCDH19 regulation of neural progenitor cell differentiation suggests asynchrony of neurogenesis as a mechanism contributing to PCDH19 Girls Clustering Epilepsy. Neurobiol Dis. 116:106–119.

Hovens, C.M., S.A. Stacker, A.C. Andres, A.G. Harpur, A. Ziemiecki, and A.F. Wilks. 1992. RYK, a receptor tyrosine kinase-related molecule with unusual kinase domain motifs. Proc Natl Acad Sci U S A. 89:11818–11822.

Huang, S.M., Y.M. Mishina, S. Liu, A. Cheung, F. Stegmeier, G.A. Michaud, O. Charlat, E. Wiellette, Y. Zhang, S. Wiessner, M. Hild, X. Shi, C.J. Wilson, C. Mickanin, V. Myer, A. Fazal, R. Tomlinson, F. Serluca, W. Shao, H. Cheng, M. Shultz, C. Rau, M. Schirle, J. Schlegl, S. Ghidelli, S. Fawell, C. Lu, D. Curtis, M.W. Kirschner, C. Lengauer, P.M. Finan, J.A. Tallarico, T. Bouwmeester, J.A. Porter, A. Bauer, and F. Cong. 2009. Tankyrase inhibition stabilizes axin and antagonizes Wnt signalling. Nature. 461:614–620.

Hulpiau, P., and F. van Roy. 2011. New insights into the evolution of metazoan cadherins. Mol Biol Evol. 28:647–657.

Hwang, W.Y., Y. Fu, D. Reyon, M.L. Maeder, S.Q. Tsai, J.D. Sander, R.T. Peterson, J.R. Yeh, and J.K. Joung. 2013. Efficient genome editing in zebrafish using a CRISPR-Cas system. Nat Biotechnol. 31:227–229.

Jao, L.E., S.R. Wente, and W. Chen. 2013. Efficient multiplex biallelic zebrafish genome editing using a CRISPR nuclease system. Proc Natl Acad Sci U S A. 110:13904–13909.

Kim, S.H., A. Yamamoto, T. Bouwmeester, E. Agius, and E.M. Robertis. 1998. The role of paraxial protocadherin in selective adhesion and cell movements of the mesoderm during Xenopus gastrulation. Development. 125:4681–4690.

Kraft, B., C.D. Berger, V. Wallkamm, H. Steinbeisser, and D. Wedlich. 2012. Wnt-11 and Fz7 reduce cell adhesion in convergent extension by sequestration of PAPC and C-cadherin. J Cell Biol. 198:695–709.

Kwan, V., B.K. Unda, and K.K. Singh. 2016. Wnt signaling networks in autism spectrum disorder and intellectual disability. J Neurodev Disord. 8:45.

Lal, D., A.K. Ruppert, H. Trucks, H. Schulz, C.G. de Kovel, D. Kasteleijn-Nolst Trenite, A.C. Sonsma, B.P. Koeleman, D. Lindhout, Y.G. Weber, H. Lerche, C. Kapser, C.J. Schankin, W.S. Kunz, R. Surges, C.E. Elger, V. Gaus, B. Schmitz, I. Helbig, H. Muhle, U. Stephani, K.M. Klein, F. Rosenow, B.A. Neubauer, E.M. Reinthaler, F. Zimprich, M. Feucht, R.S. Moller, H. Hjalgrim, P. De Jonghe, A. Suls, W. Lieb, A. Franke, K. Strauch, C. Gieger, C. Schurmann, U. Schminke, P. Nurnberg, E. Consortium, and T. Sander. 2015. Burden analysis of rare microdeletions suggests a strong impact of neurodevelopmental genes in genetic generalised epilepsies. PLoS Genet. 11:e1005226.

Leung, L.C., V. Urbancic, M.L. Baudet, A. Dwivedy, T.G. Bayley, A.C. Lee, W.A. Harris, and C.E. Holt. 2013. Coupling of NF-protocadherin signaling to axon guidance by cue-induced translation. Nat Neurosci. 16:166–173.

Lewis, J.L., J. Bonner, M. Modrell, J.W. Ragland, R.T. Moon, R.I. Dorsky, and D.W. Raible. 2004. Reiterated Wnt signaling during zebrafish neural crest development. Development. 131:1299–1308.

Li, L., B.I. Hutchins, and K. Kalil. 2009. Wnt5a induces simultaneous cortical axon outgrowth and repulsive axon guidance through distinct signaling mechanisms. J Neurosci. 29:5873–5883.

Lin, Y., X. Ge, X. Zhang, Z. Wu, K. Liu, F. Lin, C. Dai, W. Guo, and J. Li. 2018. Protocadherin-8 promotes invasion and metastasis via laminin subunit gamma2 in gastric cancer. Cancer Sci. 109:732–740.

Lu, W., V. Yamamoto, B. Ortega, and D. Baltimore. 2004. Mammalian Ryk is a Wnt coreceptor required for stimulation of neurite outgrowth. Cell. 119:97–108.

MacDonald, B.T., K. Tamai, and X. He. 2009. Wnt/beta-catenin signaling: components, mechanisms, and diseases. Dev Cell. 17:9–26.

Mah, K.M., D.W. Houston, and J.A. Weiner. 2016. The gamma-Protocadherin-C3 isoform inhibits canonical Wnt signalling by binding to and stabilizing Axin1 at the membrane. Sci Rep. 6:31665.

McAninch, D., and P. Thomas. 2014. Identification of highly conserved putative developmental enhancers bound by SOX3 in neural progenitors using ChIP-Seq. PLoS One. 9:e113361.

Morrow, E.M., S.Y. Yoo, S.W. Flavell, T.K. Kim, Y. Lin, R.S. Hill, N.M. Mukaddes, S. Balkhy, G. Gascon, A. Hashmi, S. Al-Saad, J. Ware, R.M. Joseph, R. Greenblatt, D. Gleason, J.A. Ertelt, K.A. Apse, A. Bodell, J.N. Partlow, B. Barry, H. Yao, K. Markianos, R.J. Ferland, M.E. Greenberg, and C.A. Walsh. 2008. Identifying autism loci and genes by tracing recent shared ancestry. Science. 321:218–223.

Nusse, R., and H. Clevers. 2017. Wnt/beta-Catenin Signaling, Disease, and Emerging Therapeutic Modalities. Cell. 169:985–999.

Perez-Palma, E., I. Helbig, K.M. Klein, V. Anttila, H. Horn, E.M. Reinthaler, P. Gormley, A. Ganna, A. Byrnes, K. Pernhorst, M.R. Toliat, E. Saarentaus, D.P. Howrigan, P. Hoffman, J.F. Miquel, G.V. De Ferrari, P. Nurnberg, H. Lerche, F. Zimprich, B.A. Neubauer, A.J. Becker, F. Rosenow, E. Perucca, F. Zara, Y.G. Weber, and D. Lal. 2017. Heterogeneous contribution of microdeletions in the development of common generalised and focal epilepsies. J Med Genet. 54:598–606.

Piper, M., A. Dwivedy, L. Leung, R.S. Bradley, and C.E. Holt. 2008. NF-protocadherin and TAF1 regulate retinal axon initiation and elongation in vivo. J Neurosci. 28:100–105.

Piton, A., J. Gauthier, F.F. Hamdan, R.G. Lafreniere, Y. Yang, E. Henrion, S. Laurent, A. Noreau, P. Thibodeau, L. Karemera, D. Spiegelman, F. Kuku, J. Duguay, L. Destroismaisons, P. Jolivet, M. Cote, K. Lachapelle, O. Diallo, A. Raymond, C. Marineau, N. Champagne, L. Xiong, C. Gaspar, J.B. Riviere, J. Tarabeux, P. Cossette, M.O. Krebs, J.L. Rapoport, A. Addington, L.E. Delisi, L. Mottron, R. Joober, E. Fombonne, P. Drapeau, and G.A. Rouleau. 2011. Systematic resequencing of X-chromosome synaptic genes in autism spectrum disorder and schizophrenia. Mol Psychiatry. 16:867–880.

Polakis, P. 2012. Wnt signaling in cancer. Cold Spring Harb Perspect Biol. 4.

Pologruto, T.A., B.L. Sabatini, and K. Svoboda. 2003. ScanImage: flexible software for operating laser scanning microscopes. Biomed Eng Online. 2:13.

Redies, C., N. Hertel, and C.A. Hubner. 2012. Cadherins and neuropsychiatric disorders. Brain Res. 1470:130–144.

Schindelin, J., I. Arganda-Carreras, E. Frise, V. Kaynig, M. Longair, T. Pietzsch, S. Preibisch, C. Rueden, S. Saalfeld, B. Schmid, J.Y. Tinevez, D.J. White, V. Hartenstein, K. Eliceiri, P. Tomancak, and A. Cardona. 2012. Fiji: an open-source platform for biological-image analysis. Nat Methods. 9:676–682.

Schmitt, A.M., J. Shi, A.M. Wolf, C.C. Lu, L.A. King, and Y. Zou. 2006. Wnt-Ryk signalling mediates medial-lateral retinotectal topographic mapping. Nature. 439:31–37.

Shihavuddin, A., S. Basu, E. Rexhepaj, F. Delestro, N. Menezes, S.M. Sigoillot, E. Del Nery, F. Selimi, N. Spassky, and A. Genovesio. 2017. Smooth 2D manifold extraction from 3D image stack. Nat Commun. 8:15554.

Tang, X., X. Yin, T. Xiang, H. Li, F. Li, L. Chen, and G. Ren. 2012. Protocadherin 10 is frequently downregulated by promoter methylation and functions as a tumor suppressor gene in non-small cell lung cancer. Cancer Biomark. 12:11–19.

van Roy, F. 2014. Beyond E-cadherin: roles of other cadherin superfamily members in cancer. Nat Rev Cancer. 14:121–134.

Vanhalst, K., P. Kools, K. Staes, F. van Roy, and C. Redies. 2005. delta-Protocadherins: a gene family expressed differentially in the mouse brain. Cell Mol Life Sci. 62:1247–1259.

Westerfield, M. 1995. The Zebrafish Book. University of Oregon Press, Eugene, OR.

Williams, J.S., J.Y. Hsu, C.C. Rossi, and K.B. Artinger. 2018. Requirement of zebrafish pcdh10a and pcdh10b in melanocyte precursor migration. Dev Biol.

Wu, C., L. Niu, Z. Yan, C. Wang, N. Liu, Y. Dai, P. Zhang, and R. Xu. 2015. Pcdh11x Negatively Regulates Dendritic Branching. J Mol Neurosci. 56:822–828.

Xu, Y., Z. Yang, H. Yuan, Z. Li, Y. Li, Q. Liu, and J. Chen. 2015. PCDH10 inhibits cell proliferation of multiple myeloma via the negative regulation of the Wnt/beta-catenin/BCL-9 signaling pathway. Oncol Rep. 34:747–754.

Yin, X., T. Xiang, J. Mu, H. Mao, L. Li, X. Huang, C. Li, Y. Feng, X. Luo, Y. Wei, W. Peng, G. Ren, and Q. Tao. 2016. Protocadherin 17 functions as a tumor suppressor suppressing Wnt/beta-catenin signaling and cell metastasis and is frequently methylated in breast cancer. Oncotarget.

Zhan, T., N. Rindtorff, and M. Boutros. 2017. Wnt signaling in cancer. Oncogene. 36:1461–1473.

Zhang, P., H. Wang, J. Wang, Q. Liu, Y. Wang, F. Feng, and L. Shi. 2017. Association between protocadherin 8 promoter hypermethylation and the pathological status of prostate cancer. Oncol Lett. 14:1657–1664.

Zhang, P., C. Wu, N. Liu, L. Niu, Z. Yan, Y. Feng, and R. Xu. 2014. Protocadherin 11 x regulates differentiation and proliferation of neural stem cell in vitro and in vivo. J Mol Neurosci. 54:199–210.

Zong, Z., H. Pang, R. Yu, and Y. Jiao. 2017. PCDH8 inhibits glioma cell proliferation by negatively regulating the AKT/GSK3beta/beta-catenin signaling pathway. Oncol Lett. 14:3357–3362.

